# Draxin alters laminin expression during basement membrane reorganization to control cranial neural crest EMT

**DOI:** 10.1101/407882

**Authors:** Erica J. Hutchins, Marianne E. Bronner

**Author notes:** Correspondence to Marianne E. Bronner.

## Abstract

Premigratory neural crest cells arise within the dorsal neural tube and subsequently undergo an epithelial-to-mesenchymal transition (EMT) to leave the neuroepithelium and initiate migration. Draxin is a Wnt modulator that has been shown to control the timing of cranial neural crest EMT. Here we show that this process is accompanied by three stages of remodeling of the basement membrane protein laminin, from regression to expansion and channel formation. Loss of Draxin results in blocking laminin remodeling at the regression stage, whereas ectopic maintenance of Draxin blocks remodeling at the expansion stage. The latter effect is rescued by addition of Snail2, previously shown to be downstream of Draxin. Our results demonstrate an essential function for the Wnt modulator Draxin in regulating basement membrane remodeling during cranial neural crest EMT.

**HIGHLIGHTS:**

- Cranial neural crest migrate through a laminin-rich basement membrane channel
- Perturbation of Draxin, a Wnt antagonist, alters laminin channel formation
- Draxin’s effect on laminin channel formation is largely mediated by Snail2

## 1. INTRODUCTION

Neural crest (NC) cells undergo a spatiotemporally regulated epithelial-to-mesenchymal transition (EMT) to separate from the neuroepithelium and become migratory. The process of EMT involves first breaking cell-cell and cell-basement membrane connections to detach from the neuroepithelium and delaminate (Gouignard et al., 2018; Nieto et al., 2016). The process of delamination and onset of migration involves intrinsic changes in cell polarity and cell surface proteins, including cadherins and integrins, as well as remodeling of the basement membrane and interactions with interstitial extracellular matrix (ECM) components (Christian et al., 2013; Duband, 2006; Perris, 1997). In particular, laminin is a major component of both the basement membrane and interstitial ECM that plays an essential role in regulating cranial NC migration (Coles et al., 2006; Duband and Thiery, 1987; Perris and Perissinotto, 2000).

During EMT, cells begin the delamination process by down-regulating laminin (Zeisberg and Neilson, 2009). Interestingly, members of the Snail family of transcription factors have been shown to directly repress transcription of *laminin alpha5* (Takkunen et al., 2008), which has a critical function in cranial NC migration (Coles et al., 2006). Snail genes are well-known regulators of EMT via transcriptional repression in both cancer cells and neural crest cells (Cano et al., 2000; Strobl-Mazzulla and Bronner, 2012; Taneyhill et al., 2007). Thus, there appears to be an important link between intrinsic NC EMT factors and laminin expression. However, the mechanisms underlying the interplay between Snail, laminin, and basement membrane remodeling during neural crest EMT remain unclear.

We recently characterized the role of the Wnt signaling antagonist, Draxin, in regulating the timing of onset of cranial NC EMT (Hutchins and Bronner, 2018). Interestingly, the transcription factor Snail2, the only Snail gene family member expressed within chick cranial NC (del Barrio and Nieto, 2002), emerged as a likely target downstream of a transient pulse of Wnt signaling. Given that Snail genes are known to repress expression of laminin in other contexts (Takkunen et al., 2008) and laminin is an important component of the basement membrane surrounding the neural tube during NC EMT, we asked whether Draxin may play a role in the modulation of laminin during EMT. The results show that Draxin perturbation impedes basement membrane remodeling during cranial NC EMT via changes to laminin expression/deposition. We further demonstrate that Draxin’s effect on laminin is largely mediated by Snail2. Thus, modulation of Wnt signaling via Draxin and its downstream effector Snail2 plays a critical role in regulating basement membrane remodeling during neural crest EMT.

## 2. MATERIALS AND METHODS

### 2.1 Cryosectioning and Immunohistochemistry

Chicken embryos (*Gallus gallus)* were obtained commercially and incubated at 37°C to reach the desired Hamburger-Hamilton (Hamburger and Hamilton, 1951) stage as indicated. Embryos were fixed overnight at 4°C in 4% paraformaldehyde in sodium phosphate buffer, then cryosectioned and immunostained as described (Hutchins and Bronner, 2018). Following gelatin removal, cryosections were incubated in primary antibodies overnight at 4°C and secondary antibodies for one hour at room temperature. Primary antibodies used are listed in the Key Resources Table. Species-specific secondary antibodies were labeled with Alexa Fluor 488, 568, and 647 (Invitrogen). Prior to imaging, coverslips were mounted with Fluoromount-G (SouthernBiotech).

### 2.2 Electroporations

As described previously (Hutchins and Bronner, 2018; Sauka-Spengler and Barembaum, 2008; Simoes-Costa et al., 2015), dissected embryos were electroporated *ex ovo* at Hamburger-Hamilton (HH) stage HH4. Following injection of expression constructs or morpholino oligomers, embryos were electroporated using platinum electrodes (5 pulses, 6.0V, 30ms duration at 100ms intervals), and cultured at 37°C in albumin/1% penicillin/streptomycin to HH9+.

### 2.3 Expression constructs and Morpholino Oligomers

Draxin knockdown using translation-blocking antisense morpholino-electroporation (Draxin MO, Key Resources Table; GeneTools; co-electroporated with pCIG), and Draxin overexpression using the Draxin-FLAG expression vector, were performed as described (Hutchins and Bronner, 2018). Control reagents were described previously (Hutchins and Bronner, 2018), and are listed in the Key Resources Table. In order to generate the NC1-Snail2 construct, the coding for V5-tagged Snail2 was PCR amplified from pCIG-V5-Snail2-IRES-nls-GFP (Liu et al., 2013). This V5-Snail2 fragment was then cloned into the NC1.1 M3 construct (Simoes-Costa et al., 2012), following excision of GFP by *Nhe*I and *Xba*I. All constructs and primers (Integrated DNA Technologies) are listed in the Key Resources Table.

### 2.4 Microscope image acquisition

Images were acquired using a Zeiss Imager.M2 with an ApoTome.2 module and Zen 2 Blue software, using a Zeiss Plan-Apochromat 20x objective/0.8 NA. Z-stacks were taken at 0.55µm intervals and displayed as maximum intensity projections. Images were minimally processed for brightness and contrast using Adobe Photoshop CC.

### 2.5 Statistical analysis

Statistical analyses were performed using GraphPad Prism 7. *P* values were calculated using an unpaired, two-tailed t-test and are indicated in the text. Data are presented as the percentage of the total number of sections displaying the specified phenotype (n≥8 sections among 5 embryos, per condition).

## 3. RESULTS

### 3.1 Cranial neural crest migrate through a laminin channel during EMT

In avian cranial NC, the events of EMT and neural tube closure progress simultaneously. As the neural tube undergoes separation from the non-neural ectoderm, premigratory NC cells in this region delaminate and complete EMT to migrate dorsolaterally between the newly-formed overlying dorsal ectoderm and the underlying ventral mesoderm (Baker et al., 1997; Duband, 2006; Theveneau and Mayor, 2012). The separation of the neural tube from the non-neural ectoderm occurs via a remodeling of the laminin-rich basement membrane from a single, continuous structure into distinct dorsal ectodermal and neural tube basement membranes (Duband and Thiery, 1987).

In order to correlate events of cranial NC EMT with reorganization of the basement membrane, we examined the expression of laminin (as a marker for the basement membrane) and the neural crest marker Pax7 across several developmental time points. Immunohistochemical analysis of transverse sections through wild-type chick mesencephalon revealed dynamic rearrangement of the basement membrane relative to discrete stages of cranial neural crest EMT. Between Hamburger–Hamilton stages 8+ and 9- (HH8+/9-), as specified premigratory NC cells were localized to the dorsal aspect of the neural tube, the basement membrane was remodeled to form a space between the non-neural ectoderm and neural tube, which were tightly apposed at HH8+ (Fig. 1A-B, arrowheads). At HH9, as cranial NC began delaminating from the neuroepithelium, the basement membrane at the junction of the neural tube and ectoderm remained continuous, despite expanding to encapsulate newly delaminated NC, apparently creating a barrier that blocked NC entry into the dorsolateral migratory route (Fig. 1C). By HH9+, the lateral laminin barrier disappeared to generate separate ectodermal and neural tube basement membranes. Concomitantly, cranial NC cells completed EMT and mesenchymalized to migrate away from the neural tube (Hutchins and Bronner, 2018). Because the basement membranes of the ectoderm and neural tube remained in close proximity, this separation formed a laminin-lined “channel”, through which the cranial NC migrated (Fig. 1D). Thus, during cranial NC EMT, basement membrane remodeling occurs in three successive stages, which we term “Regression” (Fig. 1B”), “Expansion” (Fig. 1C”), and “Channel Formation” (Fig. 1D”).

**Figure 1.**
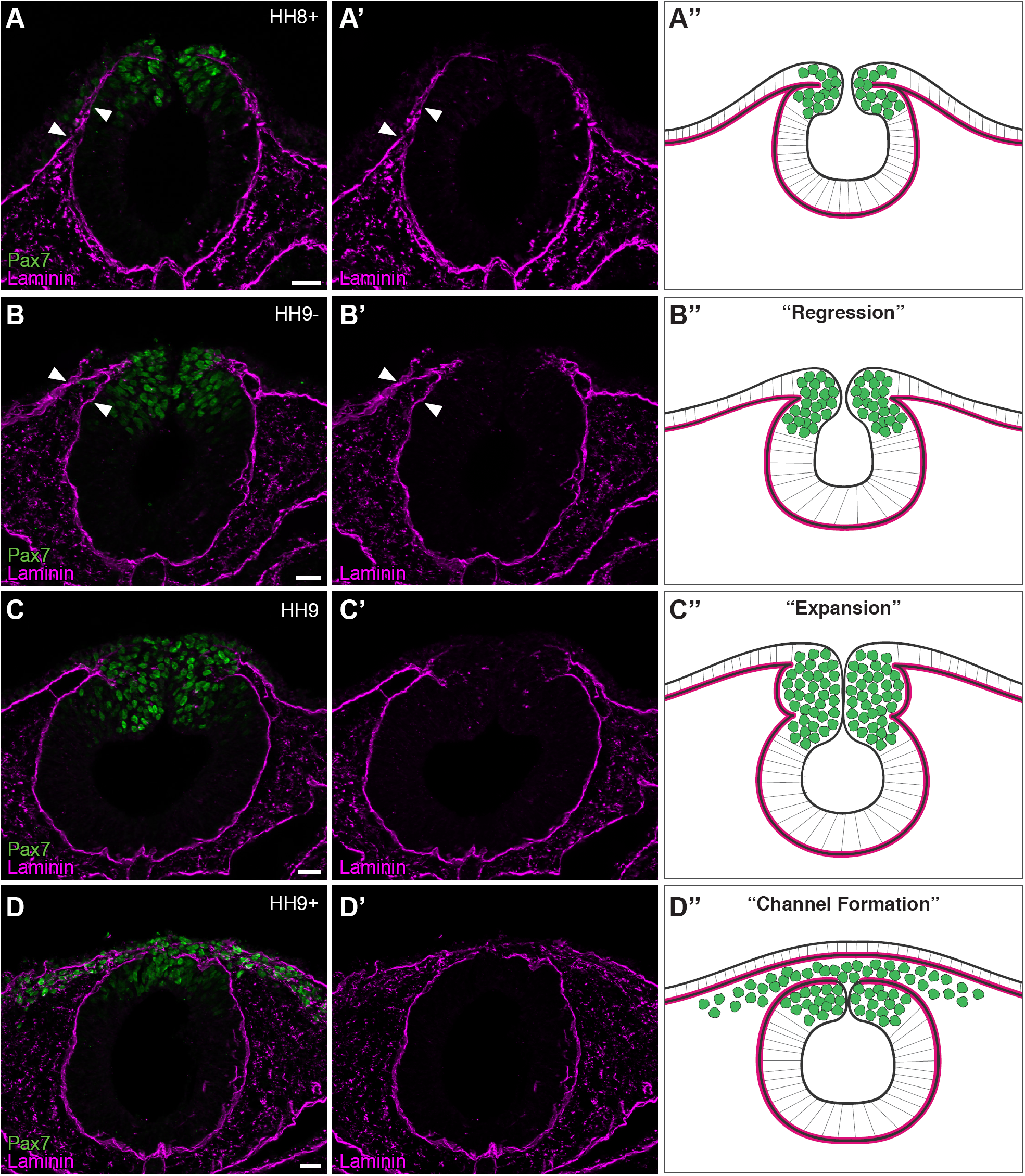
The basement membrane is remodeled during cranial neural crest EMT. (A-D) Immunostaining for Pax7 (green) and laminin (magenta) in cross sections of HH8+ (A), HH9- (B), HH9 (C), and HH9+ (D) embryos. Arrowheads in (A’) and (B’) indicate apposed and non-apposed membranes, respectively. Scale bar, 20 µm. Schematics (A”-D”) summarize immunostaining data (n=12 sections, 3 embryos per stage), where green circles represent neural crest cells and magenta lines indicate the laminin-rich basement membrane. As Pax7+ neural crest cells undergo EMT, a laminin channel forms by HH9+, creating a passage through which the cells are able to migrate.

### 3.2 Draxin knockdown impeded basement membrane remodeling during EMT

We previously identified a critical role of the Wnt antagonist, Draxin, in the regulation of cranial NC EMT; loss of Draxin induced premature NC exit from the neuroepithelium, though the cells failed to complete EMT and become migratory (Hutchins and Bronner, 2018). To determine whether Draxin played a role in basement membrane remodeling, we examined transverse sections of embryos unilaterally electroporated with either control morpholino (MO) (Fig. 2A-B) or a translation-blocking Draxin MO (Fig. 2D-E) that was previously validated (Hutchins and Bronner, 2018). Whereas control MO electroporation yielded no significant change in the incidence of laminin channel formation compared to the contralateral uninjected side (*p* = 0.63; Fig. 2C), loss of Draxin significantly inhibited laminin channel formation (*p* < 0.0001; Fig. 2F). Examination of the Draxin knockdown phenotype at stage HH9+ showed a continuous basement membrane (Fig. 2E, arrowhead), with laminin expression resembling a stage HH9-wild type phenotype (Fig. 1B). These data suggest that Draxin knockdown halts basement membrane remodeling at the regression stage. Thus, Draxin is necessary for complete basement membrane reorganization during cranial neural crest EMT.

**Figure 2.**
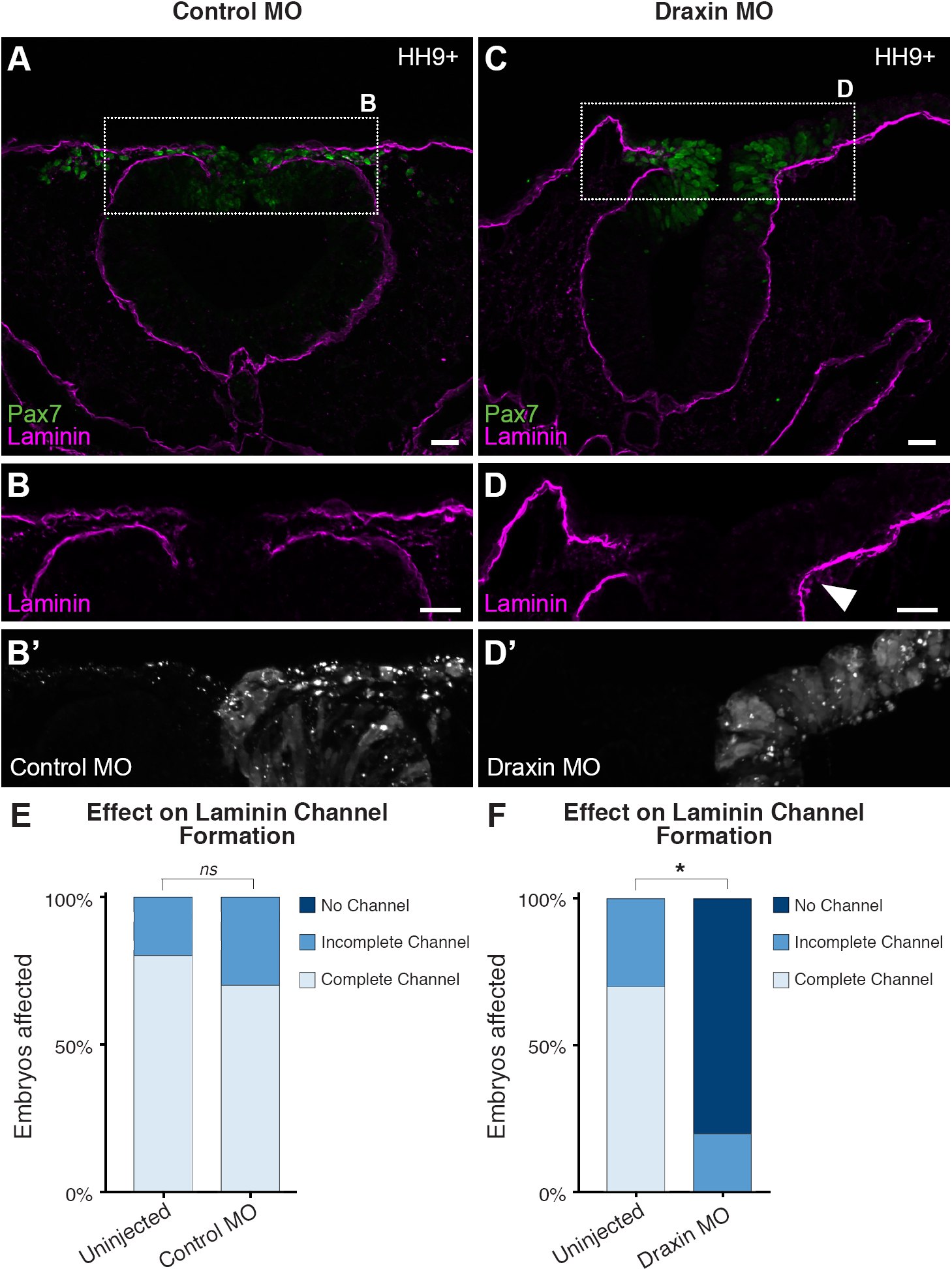
Draxin knockdown inhibits basement membrane remodeling during EMT. (A-D) Representative images for Pax7 (green) and laminin (magenta) immunostaining in embryos unilaterally electroporated with FITC-labeled control (A-B) or translation-blocking Draxin (C-D) morpholino (MO). Scale bar, 20 µm. (E-F) Quantification of the effect of MO electroporation indicated Draxin knockdown significantly reduced laminin channel formation, whereas control MO did not (*p* = 0.63). Boxes in (A,D) indicate zoomed areas in (B,E), as indicated. Arrowhead highlights area of altered laminin deposition. *, *p* < 0.0001, two-tailed *t*-test. ns, nonsignificant.

### 3.3 Ectopic maintenance of Draxin blocks laminin channel formation during EMT

*Draxin* is expressed in a transient head-to-tail wave within the premigratory NC, and its downregulation mirrors initiation of cranial NC emigration. Furthermore, ectopic overexpression of Draxin after its endogenous downregulation results in inhibited cranial NC delamination and migration (Hutchins and Bronner, 2018). Thus, we sought to assess the effects of ectopic Draxin overexpression on laminin expression during EMT. Following unilateral electroporation of pCI-H2B-RFP as a control (Fig. 3A-B) or a Draxin-FLAG overexpression construct (Hutchins and Bronner, 2018; Fig. 3D-E), we examined transverse sections of embryos for effects on laminin channel formation. Control pCI-H2B-RFP electroporations had no significant effect on laminin channel formation (*p* = 0.29; Fig. 3C). In contrast, ectopic Draxin overexpression significantly reduced the incidence of complete channel formation (*p* < 0.0001; Fig. 3F); examination of the Draxin-FLAG phenotype at stage HH9+ revealed a physical blockage of the laminin channel due to aberrant laminin expression (Fig. 3E, arrowhead), resembling a stage HH9 wild type phenotype (Fig. 1C). These data suggest that ectopic maintenance of Draxin halts basement membrane remodeling at the expansion stage, inhibiting the dissolution of the lateral laminin barrier, which creates a physical block to the migration of cranial NC away from the neural tube.

**Figure 3.**
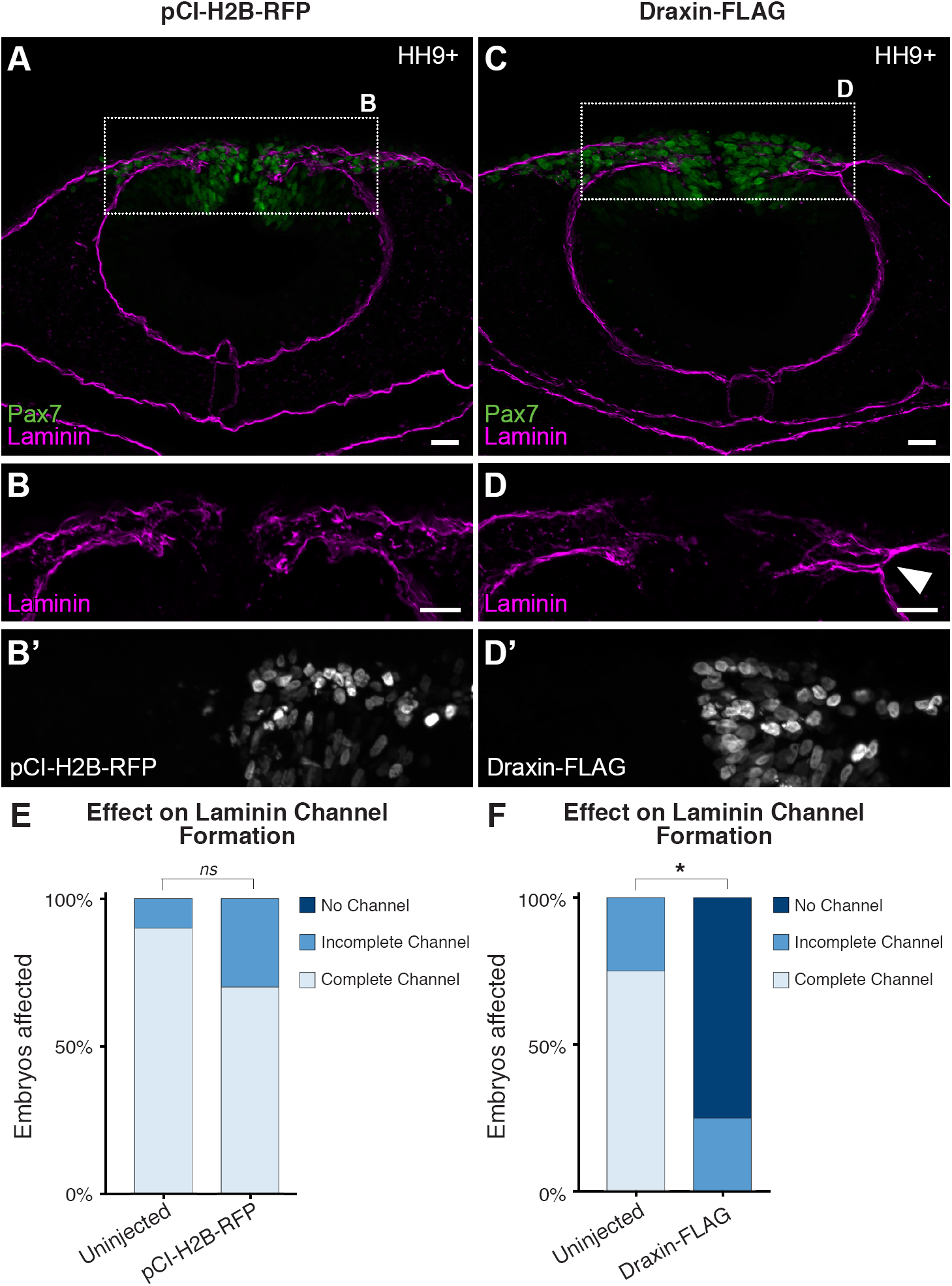
Ectopic maintenance of Draxin impedes laminin channel formation. (A-D) Representative images for Pax7 (green) and laminin (magenta) immunostaining in embryos unilaterally electroporated with pCI-H2B-RFP control (A-B) or a Draxin-FLAG overexpression construct (C-D). Scale bar, 20 µm. (E-F) Quantification of laminin channel formation indicated that, whereas pCI-H2B-RFP electroporation had no effect on channel formation (*p* = 0.29), Draxin-FLAG overexpression significantly inhibited channel completion due to altered laminin expression. Boxes in (A,D) indicate zoomed areas in (B,E), as indicated. Arrowhead highlights area of altered laminin deposition. *, *p* < 0.0001, two-tailed *t*-test. ns, nonsignificant.

### 3.4 Snail2 rescues laminin channel formation from Draxin overexpression

We have previously demonstrated that Draxin overexpression inhibited Snail2 in cranial NC (Hutchins and Bronner, 2018). Given that Snail family members have been implicated in both EMT and the transcriptional control of *laminin* (Cano et al., 2000; Takkunen et al., 2008; Zeisberg and Neilson, 2009), we asked whether the effects of Draxin on laminin channel formation and basement membrane remodeling in cranial NC are mediated by Snail2. To this end, we generated a Snail2 overexpression construct (NC1-Snail2) in which the FoxD3 NC1 enhancer (Simoes-Costa et al., 2012) drives expression of full-length chick Snail2. This construct restricts expression of Snail2 to the cranial NC following specification, in order to bypass potential deleterious effects on earlier induction and specification events. When co-electroporated with Draxin-FLAG (Fig. 4A-B), we observed complete rescue of laminin channel formation (Fig. 4C; compare to Fig. 3D-F). Together, these data suggest that Draxin controls the progression of basement membrane remodeling via modulation of Snail2.

**Figure 4.**
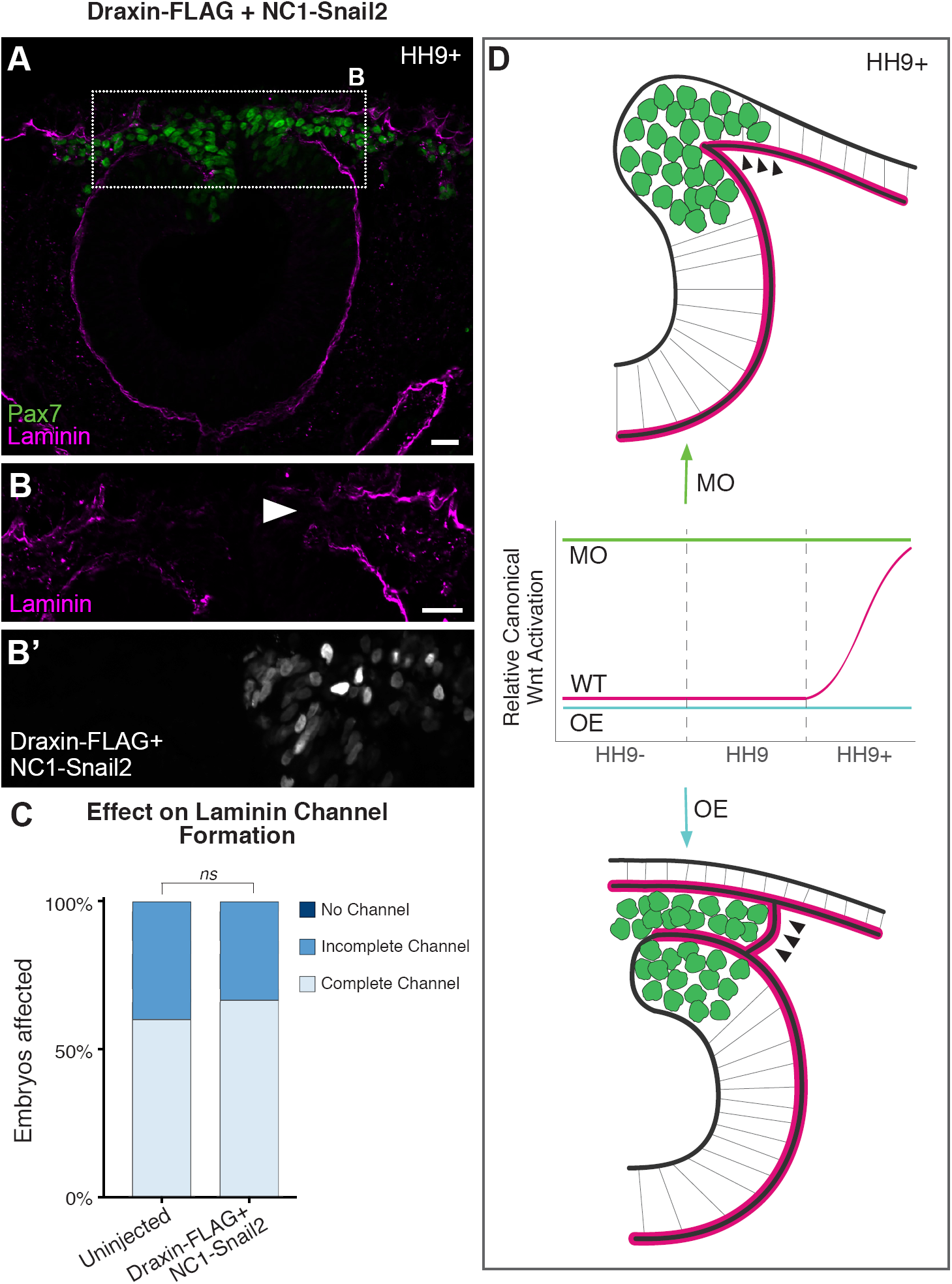
Overexpression of Snail2 rescues the effect of Draxin on laminin channel formation. (A-B) Representative images for Pax7 (green) and laminin (magenta) immunostaining in embryos unilaterally co-electroporated with NC1-Snail2 and Draxin-FLAG. Scale bar, 20 µm. Box in (A) indicates zoomed area in (B), as indicated. Arrowhead highlights rescue of channel formation. (C) Quantification of laminin channel formation indicated that NC1-Snail2 rescued the effect of Draxin-FLAG overexpression on channel formation. ns, nonsignificant (*p* = 0.53). (D) Model summarizing results and hypothesized role of canonical Wnt signaling output on basement membrane remodeling during cranial neural crest EMT. WT, wild type. MO, morpholino knockdown phenotype. OE, overexpression phenotype.

## 4. DISCUSSION

The canonical Wnt antagonist, Draxin, has critical functions in regulating the progression of cranial NC EMT. A transient pulse of *Draxin* expression tightly coordinates the events of cranial NC delamination and mesenchymalization, to allow successful migration away from the neural tube (Hutchins and Bronner, 2018). Here, we demonstrate a novel function of Draxin during EMT—in regulation of basement membrane remodeling. Our data provide a mechanistic link between Wnt signaling modulation and progressive laminin reorganization to restructure the basement membrane.

Combined with our prior study (2018), our results suggest the following model for the interplay between Snail, laminin, and basement membrane remodeling during cranial NC EMT (Fig. 4D). Under wild-type conditions, endogenous *Draxin* expression within the premigratory cranial neural crest attenuates Wnt signaling, and by extension, the Wnt target gene Snail2; reduced Wnt signaling maintains the basement membrane in a single, continuous layer that is able to progress through the regression and expansion stages. As *Draxin* is endogenously downregulated, Wnt signaling increases, allowing intrinsic NC EMT factors (*e.g.* Snail2, etc.) to dissolve the lateral laminin barrier (presumably via both transcriptional and post-translational regulation) and complete channel formation.

With loss of Draxin, Wnt signaling increases prematurely and halts laminin reorganization. We hypothesize this abrogation of basement membrane remodeling may be in part due to reduced Cadherin6B (Cad6B) expression. Cad6B is a target of proteolytic cleavage, and it was recently shown that its cleaved fragments promote laminin degradation and basement membrane remodeling during EMT (Schiffmacher et al., 2018; Schiffmacher et al., 2014). Thus, it is possible the premature reduction of Cad6B found with loss of Draxin (Hutchins and Bronner, 2018) subsequently reduces the availability of Cad6B cleavage fragments to modulate laminin. Further, because Snail1 directly represses transcription of a subtype of *laminin* during EMT (Takkunen et al., 2008), loss of Draxin may affect laminin expression in multiple ways to impede basement membrane remodeling.

With ectopic maintenance of Draxin, Wnt signaling fails to increase following NC delamination, and as such, Snail2 expression remains suppressed as Cad6B is inappropriately maintained (Hutchins and Bronner, 2018). This results in a failure to remove the laminin blockage to complete channel formation, despite an increase in Cad6B. We hypothesize that the inability to remove the laminin blockage may be due the loss of Snail2, and the resulting inability to transcriptionally repress *laminin.* This argument is strengthened by the ability of Snail2 overexpression to rescue laminin channel formation from Draxin overexpression. However, there may also be a reduction in the cleavage of Cad6B, subsequently impeding laminin degradation. Parsing whether Draxin overexpression reduces proteolytic cleavage of Cad6B may provide further mechanistic insights into the effects on laminin during EMT.

In summary, our results suggest an essential function for Draxin in regulating basement membrane remodeling during cranial NC EMT. The progressive reorganization of laminin within the basement membrane correlates with the endogenous transient pulse of *Draxin* expression. Draxin’s role as a canonical Wnt antagonist thus suggests that the events of cranial NC EMT and basement membrane remodeling are inextricably linked, and that they are tightly controlled and coordinated with canonical Wnt signaling.

## ACKNOWLEDGEMENTS

We thank Dr. M. Simoes-Costa for providing the NC1.1 M3:EGFP construct, and Dr. M. Cheung for providing the pCIG-V5-Snail2-IRES-nls-GFP construct (Addgene plasmid # 44282). This work was supported by a National Institutes of Health grant (R01DE024157 and PO1HD037105 to M.E. Bronner), and a Ruth L. Kirschstein National Research Service Award (F32DE026355 to E.J. Hutchins). The authors declare no competing interests.

## KEY RESOURCES TABLE

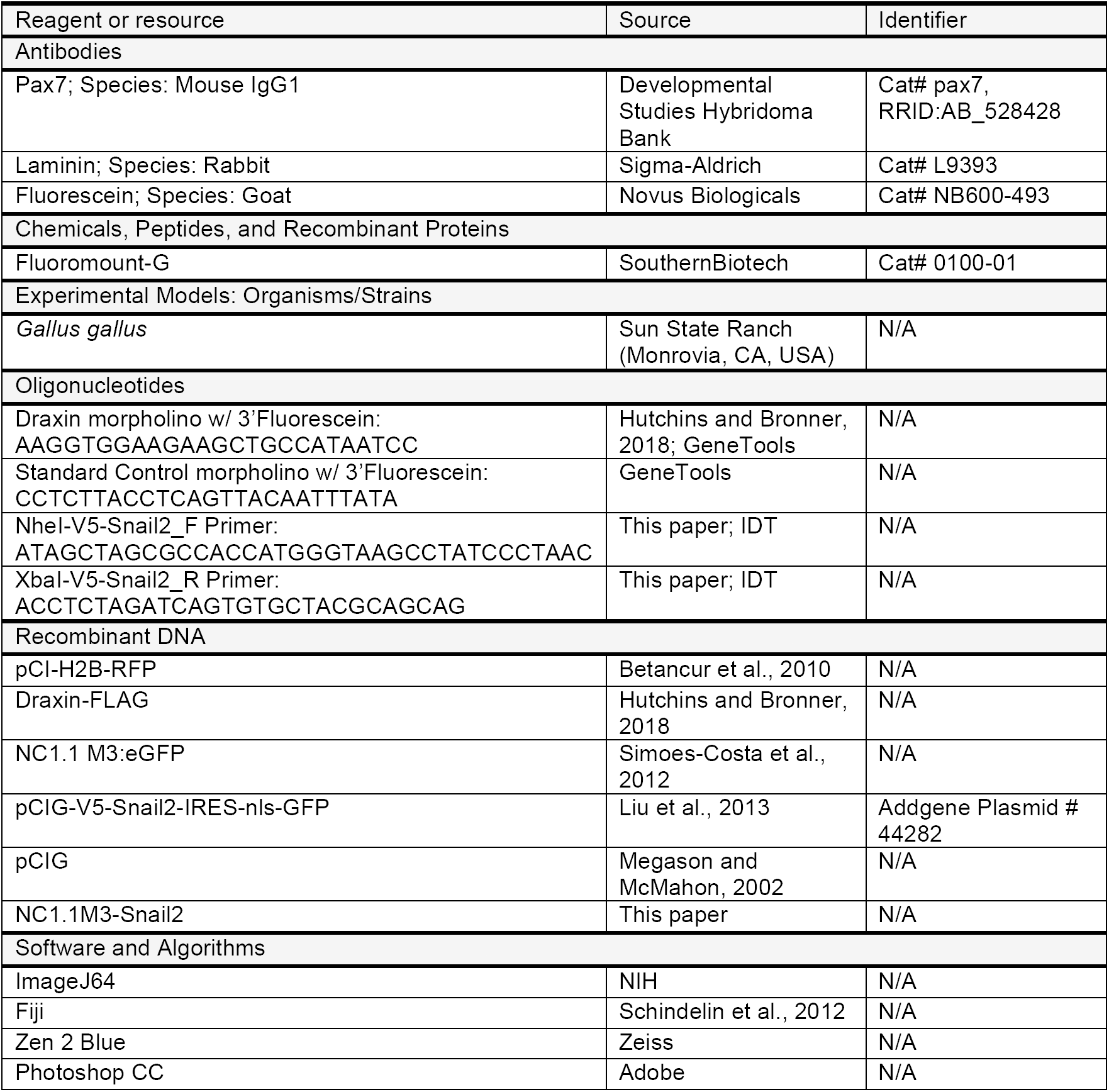

